# Drivers of vascular plant species area relationships in British broadleaved woodlands and their effects on the species area curve

**DOI:** 10.1101/2020.02.11.943282

**Authors:** Petra Guy, Simon Smart, Colin Prentice

**Author notes:** Data is available from Centre for Ecology and Hydrology, Lancaster Environment Centre, Lancaster University, Library Avenue, Bailrigg, Lancaster, LA1 4YQ, UK, https://catalogue.ceh.ac.uk/documents/ddff0f17-c95d-4415-80cb-aa9487edcb06.

## Abstract

The loss of plant biodiversity in Great Britain is a major concern, with a fifth of species endangered or vulnerable according to the latest IUCN Red List. The Government’s 25 Year Plan for the environment aims to halt this loss and build new habitats, including new woodlands. To ensure that biodiversity loss is halted in existing woodlands and gain is maximised in new ones, we need to better understand which drivers have been most influential in controlling biodiversity. Here we focus on vascular plant species’ richness.

Previous attempts to explain plant species richness have mainly focussed on alpha diversity in a consistent, fixed unit area. Here, we additionally undertake a novel analysis of the effects of environmental heterogeneity and abiotic factors on species-area relationships derived from 16 randomly placed quadrats in each of 103 semi-natural, broad-leaved woodlands across Britain. Species-area relationships were examined at two scales (4m^2^ to 200m^2^ and 200m^2^ to 3200m^2^) to explore the relationship between the drivers of species richness and the exponent z, of the canonical species-area curve, S = cA^z^. We also explore the use of a new metric ζ_r_, based on zeta diversity to quantify heterogeneity. Zeta diversity quantifies the number of species shared between multiple combinations of plots.

Habitat heterogeneity increased species richness, as did the proximity of the woodlands to surrounding natural habitats. Higher levels of soil organic matter and the progression of woodlands to later successional stages, decreased species richness. Richness was also seen to have a unimodal response to soil acidity with a peak around pH 6. At the smaller scale, heterogeneity elements in the woodland such as riparian zones or coppicing led to an increase in the value of the exponent of the species area curve. At the larger scale, species turnover led to an increase in the exponent of the curves while succession led to a decrease. At both scales, soil organic matter content had a negative effect. ζ_r_ was found to be a significant and important variable and to affect both species richness and the slope of the species accumulation curves at larger scales.

**Synthesis:** Habitat heterogeneity measures included the presence of coppicing, open areas such as rides and riparian zones and the difference between species assemblages in different plots in the woodland. Results suggest that to maximize vascular plant diversity, woodlands should be managed for heterogeneity. In addition, the increase in richness with exposure to surrounding natural habitats suggests that woodlands benefit from being embedded in more benign habitats and further, that land management surrounding woodlands has a clear role to play in supporting biodiversity within woodlands. This is an area were Agri-environment schemes have an important role.

## Introduction

Biodiversity worldwide continues to decline (Butchart et al., 2010x, Tittensor et al., 2014) and Britain is no exception to this trend (Hayhow et al., 2016). The IUCN Red List for the UK shows 22% of plant taxa as critically endangered, endangered or vulnerable (Stroh et al., 2014). Biodiversity is important for ecosystem function (Gaston & Spicer, 2004), with woodlands being especially important in supporting ecosystem functioning and supply of ecosystem services. These include improvement of water quality, flood control (McCulloch and Robinson, 1993) carbon sequestration (Brainard, Bateman and Lovett, 2009; Ostle *et al*., 2009)nectar provision within farmed landscapes (Baude et al 2016) and the provision of habitats for other taxa (Amar et al., 2006, Fuller & Warren, 1993).

The aims of the new 25-Year Environment Plan (Defra, 2018) include achieving “thriving plants and wildlife”, planting 180,000 hectares of new woodland and ensuring that existing woodlands are better managed. However, more woodland is not necessarily the answer to biodiversity loss if we cannot ascertain why our current woodlands are not thriving.

This study was aimed at finding the main drivers of vascular plant species richness in British broad-leaved semi-natural woodlands to provide research-based evidence on how best to manage, maintain and increase their plant biodiversity. We analyse diversity as the slope of the species area relationship expressed across multiple samples at the national scale.

### SUMMARY OF FACTORS THAT HAVE BEEN SHOWN TO AFFECT SPECIES RICHNESS

Many factors have been shown to affect vascular plant species richness in woodlands, including climate, edaphic properties, eutrophication, shading, presence of invasive or dominant species, habitat fragmentation and isolation, disturbance and previous land use, to name but a few. (Dzwonko and Loster, 1988; Dumortier *et al*., 2002; Dzwonko and Gawroński, 2002; Petit *et al*., 2004; Kirby *et al*., 2005; Brudvig and Damschen, 2011; Hill and Preston, 2015). Rather than carry out an in-depth review of all these effects, we review those factors whose effects we can represent from the CEH Woodland Survey database (Kirby et al., 1971) and justify the selection of these predictor variables by citing support in the published literature. This is important due to the model selection and averaging procedure used as part of this analysis which requires justification for the inclusion of variables from prior knowledge.

Studies of beech and hornbeam woodlands (Kooijman and Cammeraat, 2010) report lower species richness at lower soil pH and when soil organic matter content is higher. This was true for both taxa, suggesting that the effect was not the result of the physical barrier created by slowly degrading beech litter.

Cornwell & Grubb (2003) showed that the greatest richness in woodlands occurred on nutrient rich soils, reporting a unimodal response with a peak around Ellenberg N of 7 for shade loving species. Since most macronutrients are available around pH5/6 this implies that a unimodal richness response in woodlands should be expected.

Ohlemuuler & Wilson, (2000) showed a decrease in richness with latitude in New Zealand for woody species, but this variation was not seen in herbaceous species. In studies on forests stretching from the equator to 60° latitude, Gillmann (2014) found that net primary production increases toward the equator and concludes that, since vascular plant richness is correlated with net primary production, richness will also increase. However, this effect may not be evident given the modest change in latitude in our UK data.

The 2010 Lawton report (Lawton et al, 2010) summarised the need for more connectivity within our landscape. Many authors, in many different settings, have shown that fragmentation reduces biodiversity. Brudvig et al (2009) showed that plants which depend on animals for dispersal increased when corridors connected experimental patches of regenerating pine forest. Thiele et al., (2018) studied plant species richness in agricultural, fragmented landscapes and concluded that connectivity increased the richness of grassland and wetland plants. Hannay et al. (1997) used the amount of forested area around a woodland to quantify isolation, and isolation and showed that richness decreased as the amount of surrounding woodland decreased. Petit et al., (2004) showed that fragmentation constrains the dispersal of ancient woodland species in British lowland woodlands. Dzwonko & Loster (1988) showed that woods which that had been isolated for longer in the agricultural landscapes in the Carpathian foothills had lower species richness.

Habitat heterogeneity is known to be important for biodiversity (Gardener, 2010). It has been shown to increase richness when it takes the form of woodland management (Boch et al, 2013, Schmidt, 2005), presence of less shaded features such as gaps, rides and paths, number of soil types, the number of different habitats the woodland contains (Honnay et al, 1999) and windthrow (Smart et al, 2014).

### SPECIES AREA CURVES AND DRIVERS AND z VALUES

Species area curves describe the change in the number of species accumulated over an area. They can be used to predict numbers of species over areas larger than those at which sampling is possible or to see if sampling is exhaustive. These curves are often described by the power model,

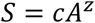

Where S is the number of species in area A, c is the number of species in a unit area and the exponent z describes how the slope of the curve changes with area. There have been many debates about the biological meaning of the z value. Z values of around 0.25 were first proposed by Preston (1960) as being the result of sampling from log-normal species distributions. This value would be expected to occur when sampling from similar habitats, for example from nested quadrats. Other authors have suggested that the z value has no direct biological meaning (Connor et al, 1983, Tjorve & Tjorve, 2017) but is a result of species distribution and aggregation. Using neutral models, Cencini et al (2012) predicted that population size affects the shape of the curve while Sugihara (1979) concludes that the value of 0.25 reflects community structure.

In this study, species were accumulated in different ways depending on the scale. At the smaller scale, species area curves were generated for species across nested areas. These are referred to as Type I curves (Scheiner, 2003). At this scale a z value of around 0.25 is expected. At larger scales we also accumulated species across all the non-contiguous quadrats in the site. These curves are referred to as species accumulation curves and at this scale habitat heterogeneity is likely to exert a greater influence on the z value (Storch et al, 2014). In similar work, Shen et al (2009) considered the effect of tree density, soil nutrients, pH, slope, elevation and aspect and heterogeneity and concluded that dispersal and habitat heterogeneity jointly are required to explain the shape of species area curves. Baldi & Sadler (2017) concluded that habitat heterogeneity, as quantified by the number of habitat types as shown through CORINE land cover maps, was more important than area for expressing species richness.

In this work, the combined influence of local abiotic factors and woodland heterogeneity on the exponent of curves constructed over these two different scales, was examined.

### HABITAT HETEROGENEITY – THE USE OF ZETA DIVERSITY

Habitat heterogeneity is known to have a major influence on biodiversity (Gardener, 2010), but it is not trivial to quantify and has been expressed in many ways. Honnay et al., (1999) used length of rides, number of soil types and a ratio of perimeter and area. Schmidt (2005) used the presence of logging trails, and Bàldi (2008) used the number of CORINE land cover codes. Here we quantified heterogeneity by combining several variables (Table 2) into a Heterogeneity Index (HI). The index has some shortcomings since it does not enumerate the numbers of each feature that occur in the woodland but merely records a feature as present irrespective of quantity or extent. This will reduce the sensitivity of the index. In addition, the number of elements identified and will depend on how exhaustively the entire site was surveyed.

A new metric was also generated based on zeta diversity (Hui & McGeoch, 2014) to provide a simple, objective measure of heterogeneity in species composition between plots within each woodland.

### ZETA DIVERSITY

Zeta diversity is the average number of species shared between multiple plots, (Hui & McGeoch 2014) and can be used to show compositional change in species assemblages, (McGeogh et al., 2017) and model drivers of species turnover, (Latombe et al., 2017).

As an example, consider four plots with sets of species S_1_, S_2_, S_3_, S_4_. Let S_i_ represent the set and number of species in plot 1. Four orders of zeta can be calculated in the following way.

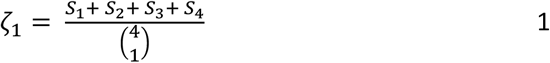

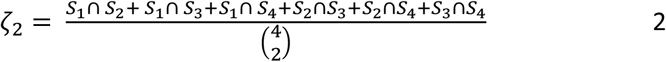

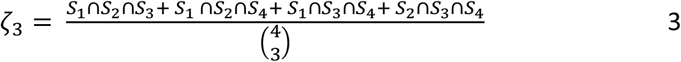

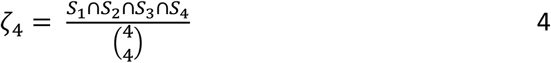

S_i_∩S_j_ is the number of species shared between plots i and j. For a woodland with 16 plots, zeta can be calculated to ζ_16_ as shown in figure 1.

**Figure 1.**
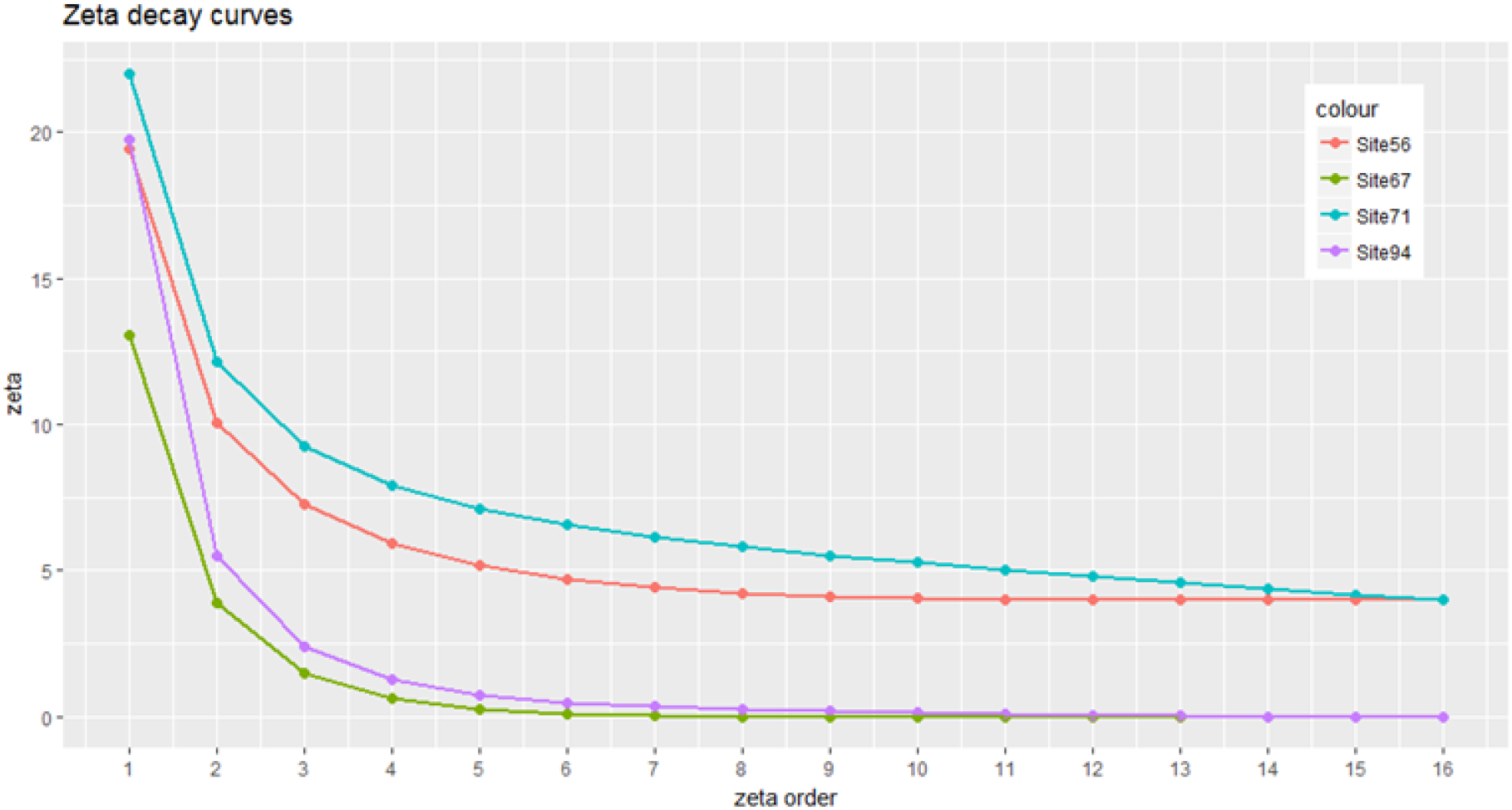
Examples of zeta decay curves calculated for 4 sites in the Bunce survey. The gradient of the initial part of the curve was used to create a metric and used as an explanatory variable to represent species heterogeneity in the woodland.

ζ_1_ is the average number of species in each plot and is therefore equivalent to mean α diversity. ζ_2_, is the average number of species shared between every combination of two plots.

The slope of the zeta decay curve varies with the probability of finding new species in new plots (Hui & McGeogh, 2014) and the number of common and rare species (Latombe et al., 2017). If plots contained the same species, then ζ_1_ - ζ_2_, = 0 and the initial gradient of the curve = 0. If all plots contained different species then ζ_2_ = 0, and the initial gradient of the curve would equal ζ_1_.

A homogeneous woodland is more likely to contain species that occur in many plots, whereas a heterogeneous woodland is more likely to have plots containing different species. This suggests a method of comparing heterogeneity between woodlands using the ratio,

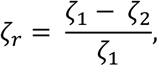

ζ_r_ is then the initial gradient of the zeta decay curve normalised by ζ_1_ to give a value between 0 and 1. The more species-compositionally homogeneous the woodland, the closer the value of ζ_r_ to zero. The zeta values for each woodland were calculated using the zetadiv function in R, (Latombe et al., 2018). The ζ_r_ coefficient was calculated and added to the set of predictor variables.

## In summary, we sought to answer three questions

1. Which variables had most influence on species richness of the entire woodland?
2. Which variables had most influence on the exponent of the species area curve and did this change with scale?
3. Can the zeta diversity metric ζr be used to represent compositional heterogeneity?

## Materials and methods

### DRIVERS OF VASCULAR PLANT SPECIES RICHNESS ACROSS ENTIRE WOODLANDS AND AT TWO DIFFERENT SCALES

The drivers of vascular plant species richness were studied in two ways. Firstly, by examining the effect of a range of variables on the species richness of a fixed area within 103 different woodlands and, secondly, by looking at the scale dependence of species richness as expressed in the canonical species area curve (SAC), *S = cA^z^*, and the effect of habitat heterogeneity and environmental conditions on the exponent, *z* of these curves. The same modelling was applied to both tasks, that is, species richness and the z values at the two different scales were used as response variables across the same set of explanatory variables.

### DATA

We used a dataset extracted from the UK Centre for Ecology and Hydrology’s woodland survey database (Wood et al 2017). One hundred and three woodlands across Great Britain were surveyed between 2000 and 2003. Sixteen square plots were randomly located in each woodland, each having a total area of 200m^2^. Further, each plot was itself nested with sub-areas of 4m^2^, 25m^2^, 50m^2^, 100m^2^ and 200m^2^.

All vascular plants and a restricted list of bryophytes were recorded within each plot by searching each concentric nest starting at the central nest and then working outwards only recording extra species recorded in each larger nest. Additional plot data were collected, including diameter at breast height (DBH) of trees, live basal area (LBA) of trees and shrubs, soil pH, soil organic matter (SOM) content, National Vegetation Classification (NVC) code, and major soil group (MSG). Additional information was also recorded where pre-specified lists of woodland attributes and features occurred in each plot. These included signs of woodland management, the presence of riparian zones and open areas such as glades or rides. Table 1 summarises the information taken from the dataset and details how it was transformed to create predictor variables.

**Table 1.** *Summary of predictors created from the variables in the dataset. Some plot scale variables, such as pH, were averaged across the woodland to give a site level variable*.

| Variable | Description |
| --- | --- |
| Heterogeneity Index (HI) | The presence of features that diversify the woodland were recorded where present. These included ponds, ditches, buildings, rock exposures and dead wood. The count of recorded features was used as an explanatory variable - HI. The greater the HI, the more elements within the woodland that could increase richness by increasing heterogeneity of conditions. |
| Northing | Latitude |
| Buffer | The proportion of land around the wood (to 3500m from the edge) that is not arable, improved grassland or urban. The larger the buffer, the more the surrounding area can be considered as a benign mosaic of habitats assumed to buffer the woodland against loss of typical woodland species and incursion of nutrient-demanding, ruderal species. This variable therefore equates to the inverse of isolation and measures the favourability of the matrix within which each woodland was embedded. |
| Number of major soil groups (MSG) | The number of major soil group codes in each woodland. This is an alternative indicator of woodland heterogeneity. |
| Number of National Vegetation Classification (NVC) codes | The number of different NVC codes in each woodland. The NVC code represents the assignment of the plant species assemblage to a phyto-sociological classification of British plant communities. The number of different NVC codes therefore describes the number of different plant assemblages present. This is equivalent to the number of CORINE land cover codes used by Baldi (2008). Although the NVC code might correlate with richness, the number of codes will not, since it is possible that each code could represent a low richness assemblage. However, since NVC codes represent a different species assemblage with different abiotic factors, if there are more codes, it is more likely that the woodland has greater heterogeneity. |
| pH | The pH of a 15 x 5cm soil sample from the centre of each plot |
| Soil Organic Matter (SOM) content | The SOM content |
| Live Basal Area (LBA) | The LBA of all saplings, trees and shrubs recorded in a plot. This variable is a measure of shading. |
| Diameter at breast height (DBH) | The DBH of all trees recorded in each plot. As with LBA, this variable is a proxy for shading. |
| Tree Density (TD) | Count of trees per plot. Again, tree density is an alternative indicator for shading. |
| Area ratio | The ratio of area to perimeter of the site. This is a measure of the number of habitats the woodland is exposed to (Honnay et al, 1999). |
| $\zeta_r$ | The ratio of $\zeta_1$-$\zeta_2$ to $\zeta_1$, quantifying species turnover between plots. |

**Table 2.** Most influential variables from the gradient boosted machine

| Parameter | Relative influence |
| --- | --- |
| $\zeta$ | 16.37 |
| Northing | 13.25 |
| meanLBA | 11.42 |
| PHI | 10.88 |
| mean pH | 9.92 |

A sample-based estimate of the λ diversity of each woodland was calculated to be the unique number of species in the total 16 plots surveyed. The variables within the database were at different scales, plot level and woodland level. Plot level variables, such as pH, DBH or SOM, were scaled appropriately to the richness by taking the mean per woodland. Site level variables were buffer, number of NVC codes, number of major soil groups, area ratio and HI

### STATISTICAL MODELLING

Two statistical modelling approaches were taken to address the first question. First, model selection using Akaike Information Criteria followed by model averaging (Burnham and Anderson, 2004) and second, a gradient boosting machine (GBM) (Natekin and Knoll, 2013). Both of these approaches were used to examine which of the predictor variables had the greatest effect on the species richness of the same area, 3200m^2^ i.e. across the 16 x 200m^2^ plots in each of the 103 woodlands.

The model selection procedure only was then repeated using the exponent, z, of the species area curves as the response with the same set of predictor variables over two scales; 4m^2^-200m^2^ (within-plot level) and 200 – 3200m^2^ (between-plot level). Finally, we evaluated the importance of ζr as a heterogeneity measure by observing the behaviour of this predictor in the above models.

### MODEL SELECTION AND AVERAGING

Model selection followed by model averaging is an ideal method for isolating variables which are most important for predicting a response when the relationships are linear, (Kimberley et al, 2014; Symonds & Moussalli, 2010). The process is frequently used in ecology and has been described in detail elsewhere (Burnham and Anderson, 2004; Johnson and Omland, 2004; Wagenmakers and Farrell, 2004; Zuur, Ieno and Elphick, 2010; Symonds and Moussalli, 2011). Therefore, a brief synopsis only of the steps involved will be given here, and is summarised in figure 2.

**Figure 2.**
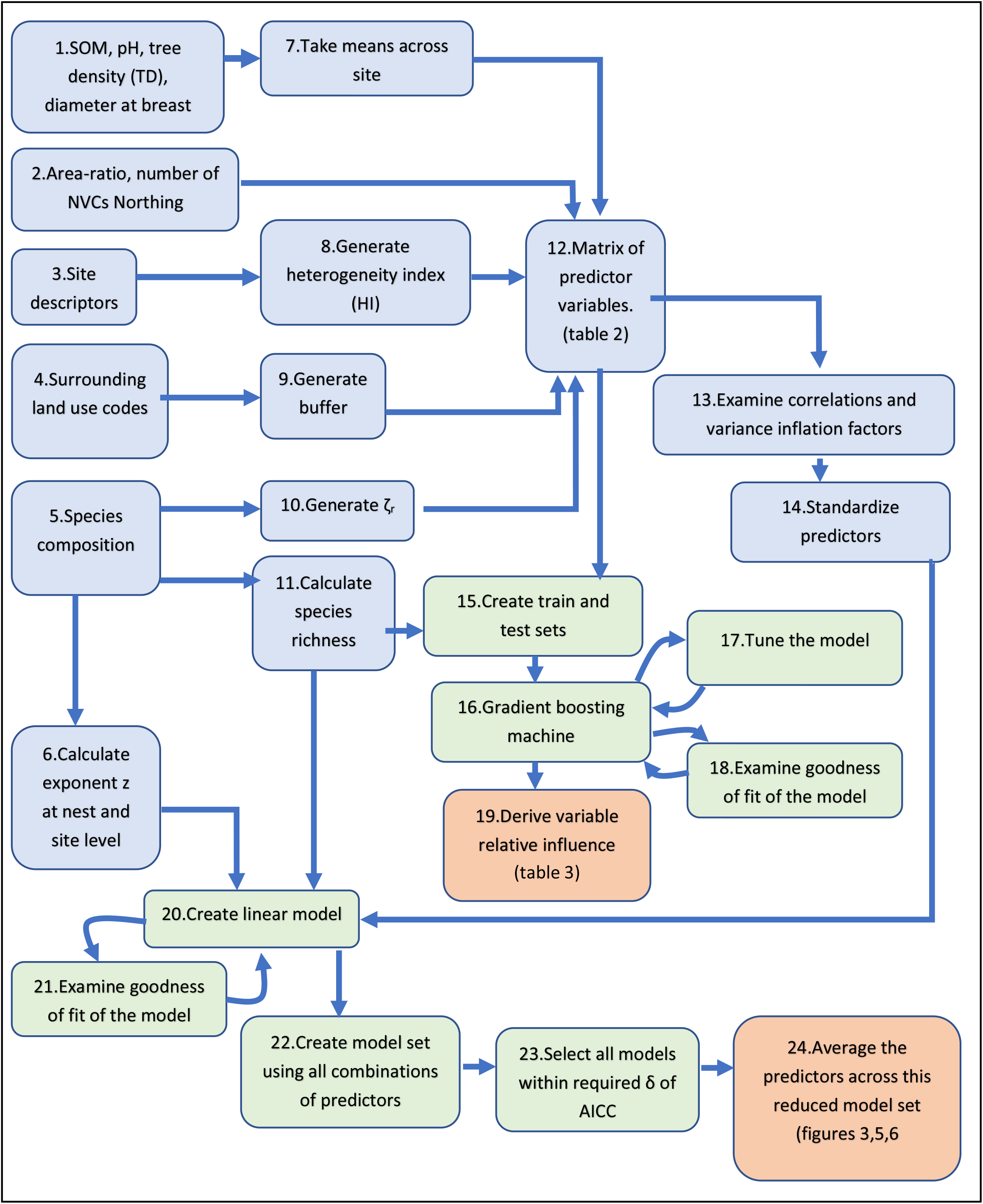
Steps involved in modelling species richness and z slopes using model averaging and GBM. Blue = model inputs, Green = quantitative modelling steps, Orange = model outputs. This diagram summarises the standard processes followed when carrying out statistical modelling using model selection and model averaging or decision trees.

Firstly, predictor variables were chosen which are known to have biological significance, (see table 1, steps 1-4, 7-9,12 in figure 2). This is because model selection is a relative measure. Including predictors with low predictive power will result in a poor set of models. This method will always produce a “best” model set across which predicators are averaged, but that model set is only “best” relative to all the other models generated. Thus, a poor set of predictors will be unable to generate a set of models with good predictive power, (Symonds and Moussalli, 2011). This means that standard data exploration protocols for linear models was followed, such as examining collinearity between predictors (step 19, figure 2) and considering R^2^ of the initial linear model (step 21 figure 2), (Zuur, Ieno and Elphick, 2010; Freckleton, 2011)). In this case we found that all variables had Spearman and Pearson correlation coefficients below 0.54. The variance inflation factors of the models were also examined and were all below 2. None of the models showed any pattern in the residuals. For the species richness models R^2^ was 0.47. For nest z, 0.33 and for plot z, 0.47.

Once the predictor variables were selected the dataset was standardized (step 14 figure 2) as suggested by Gelman, (2008). The input predictors are standardized such that two standard deviations of change in the predictor variable will result in the effect size change in the response. A linear model was created using all the variables (step 20 figure 2). The dredge function in the MuMin package in R (Barton, 2018) was used to create a set of models containing all permutations of the variable set (step 22 figure 2). These were ranked according to the AIC (step 23 figure 2). The model with the lowest AIC is considered the model that best represents the data.

All models within an AIC of between two to seven can be considered as equally good (Burnham et al., 2010) and used to create a model set. The parameter effect sizes are then averaged across this model set and a 95% confidence interval obtained (step 24 figure 2). If this confidence interval does not cross zero, these predictors are considered significant, (Kimberley et al., 2014). In this work, models within AIC of 2 (δ < 2) of the highest ranked model were initially selected. The effect of increasing δ was then examined. If more variables were introduced which had confidence intervals that crossed zero, the lower value was used to avoid including redundant models (Grueber et al., 2011). The relative importance of each effect size was also obtained, (Burnham, 2015). Relative importance is a measure of the probability that a variable is present in the model set, so the greater the importance of a parameter, the more likely it is to be found in a model. A relative importance of 1 means that a variable appears in every model in the model set.

### GRADIENT BOOSTING MACHINES

In addition to model selection and model averaging, a gradient boosting machine was used. The benefit of using these two modelling methods is their ability to detect different responses. Model-averaging detects linear relationships. Decision tree methods such as GBMs may not be as successful at detecting linear relationships, but they can detect more complex patterns in the data, (James et al, 2013).

Again, the strengths of this method and the procedure for its application have been examined and described by other authors (Ath, 2007; Elith, Leathwick and Hastie, 2008; Natekin and Knoll, 2013)) and a brief summary will therefore only be given here.

Gradient Boosting Machines (GBMs) are ensemble decision tree algorithms where recursive binary splits divide the data into regions, such that the root mean square error (rmse) of the response variable is minimized, (James et al., 2013). In addition, GBMs weight the observations such that those which were difficult to classify after a split are given increased weight. The next tree is then generated using these weighted observations. GBMs are suited to analysing complex ecological datasets because they can fit non-linear relationships and are not sensitive to scale, (De’Ath & Fabricius, 2000; Elith et al., 2008). They have been shown to be successful, for example, in finding environmental variables which explain species richness of vertebrate groups (Mouchet et al., 2015). In this analysis the gbm package in R was used, (Ridgeway, 2017).

The data was split into train and test sets using a 75/25 ratio. The GBM also uses a random subset of training data and so automatically provides a validation set. The values that are not selected can be used to give an “out of bag” (OOB) prediction error for the model.

Hyper parameters define the structure of the GBM and must be chosen to optimise the model performance. This was achieved through a hyper parameter tuning grid; a matrix containing a set of permutations of hyper parameters. The model was run across the hyper parameter tuning grid and the hyper parameters which minimized the OOB error were used in the final model.

Over fitting was addressed using the ntrees function of the gbm library. This function looks at the prediction error on the OOB set, and the number of trees is set as the value when this error stops decreasing and starts to increase. The rmse error of the train was half that of the test set, suggesting some overfitting. However, the test set rmse was comparable to that obtained using model-averaging.

To extract the variables which most influence species richness the relative influence (RI) of each variable was used, (Natekin & Knoll, 2013). The percentage decrease in rmse is averaged for each variable over each time a variable was chosen to make a binary split in the data. The variable which has greatest RI is best at splitting the data and predicting the response.

Decision tree methods such as GBMs sometimes do not give stable results for RI, (Nicodemus et al., 2007). Therefore, each model was repeated 100 times and the variables ranked by the average values of RI. The value of RI is relative and therefore subjective. In this work variables which had RI above 10% were selected as important for species richness.

### SCALE DEPENDENCE OF SPECIES RICHNESS – SPECIES AREA CURVES

We calculated species area curves across the nests from 4m^2^ to 200m^2^ and across sites, from 200m^2^ to 3200m^2^. The former curves correspond to type I nested curves as described by Scheiner (2003). In order to derive z at this smaller scale the species were accumulated across the five nests in each woodland with plot as a random effect.

At the larger scale, species were accumulated across the plots from 200m^2^ to 3200m^2^, as shown in figure 3. At this scale, species can be accumulated in 16! alternative ways i.e. every sequential arrangement of each of the 16 plots. The average of a subset of these accumulation curves is often used to calculate the slope of the species area curve. However, there are potential problems with this method when the area over which the accumulation occurs is heterogeneous, and spatial autocorrelation should be taken into account, (Bacaro et al, 2012; Bevilacqua et al, 2018, Chiarucci et al, 2009). This averaging method is effectively selecting all species for accumulation as if they have equal probabilities of occurring in any plot in the woodland, i.e. the woodland is homogeneous. This is not the case for this data, the species are grouped into assemblages. Therefore, we took a different approach to maintain this spatial information.

**Figure 3.**
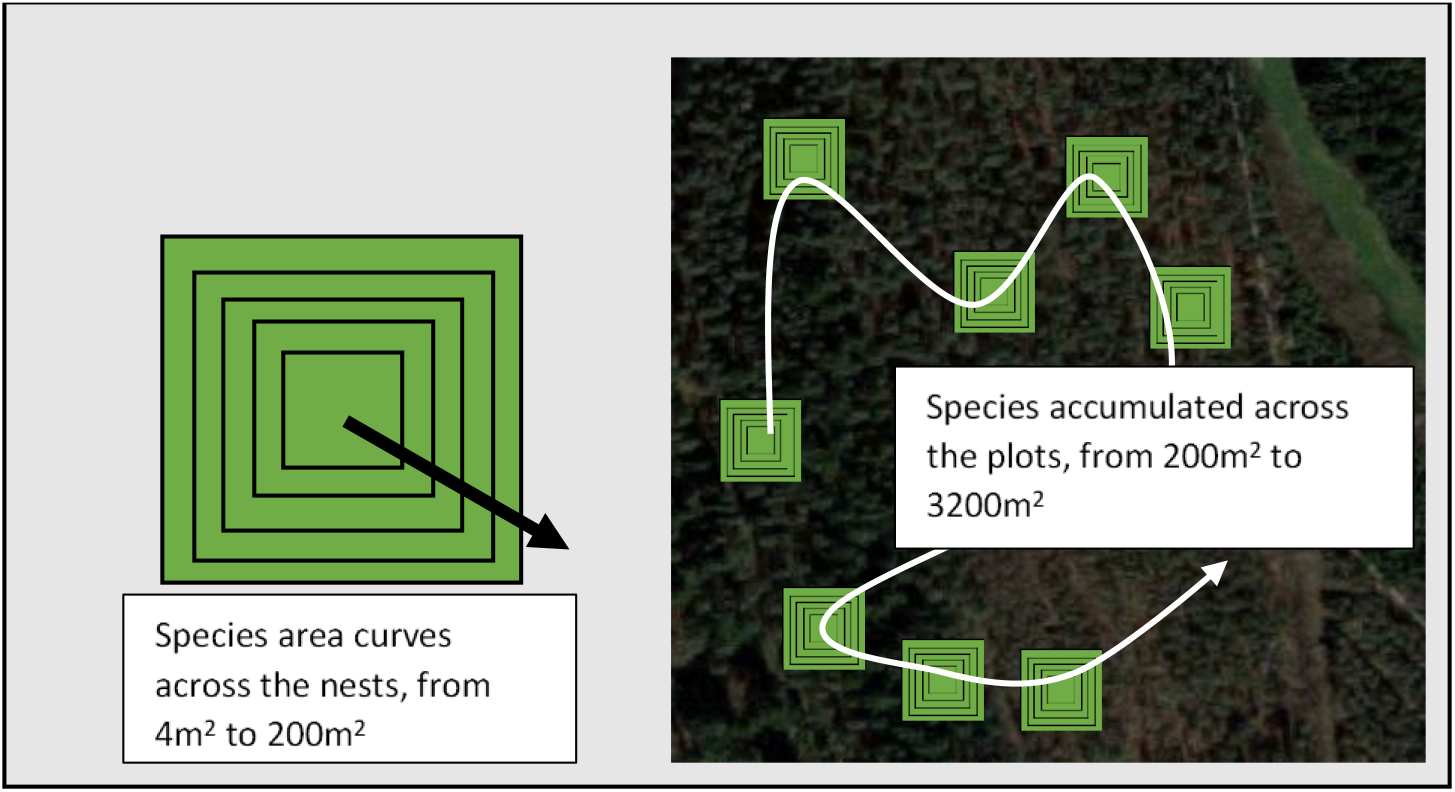
Two scales of species area dependence. At the within-plot scale species area curves were calculated for areas from 4m^2^to 200m^2^. At the between-plot scale species were accumulated across the 16 plots such that area increased from 200m^2^ to 3200m^2^.

To overcome this in a simple to execute algorithm, two curves were created. First, species were accumulated in such a way as to give a minimum gradient to the species accumulation curve. To do this, the richest plot is selected as a starting plot. The plot which shares the most species with the starting plot is accumulated next, and so on. This is equivalent to grouping all similar species assemblages together. Secondly, species were accumulated to give a maximum gradient. For this curve, the most species poor plot is selected first. The second plot to be accumulated is the one that has the least number of shared species with the starting plot. This is equivalent to arranging the plots in the most heterogeneous manner. Since these two cases represent two extremes in the woodland (the most homogenous and the most heterogeneous) an average was taken of the two resulting curves. Whilst this average does have the effect of homogenising the woodlands to some extent, we expect that this is not as extreme as averaging over all possible accumulation curves and will preserve some spatial heterogeneity.

This spatially explicit method is similar to a type III curve, as described by Scheiner (2003), where mean diversities of adjacent plots are calculated, but here adjacency is in terms of species assemblage’s similarity or dissimilarity. Using this method to fit log(S) to log(A) we found that 94% of the variance in log(S) was explained for three quarters of the woodlands.

At both scales the slope parameters of the log/log transform of the curves were used as the response variable z.

These linearized slope coefficients from the species area curves were then subject to explanatory modelling with the same variables used to model species richness, as listed in table 1.

## Results

### SPECIES RICHNESS OF FIXED AREA OF 3200M^2^

Figure 4 shows the model-averaged effect sizes and 95% confidence intervals for the total richness (λ diversity) accumulated across all the 16 200m^2^ plots in each woodland. Area of semi-natural habitat around each site (Buffer), number of NVC codes (no_NVC), Heterogenity Index (HI) and ζ_r_ all had a significant positive effect on species richness with similar effect sizes. Mean diameter at breast height (meandbh) and mean SOM had a negative effect, again, with similar effect size. All significant variables were included in every model in the model set except ζ_r_ which shows an 80% probability of inclusion. The effect sizes of all significant variables were similar, with no one parameter clearly having a greater influence on richness than any other.

**Figure 4.**
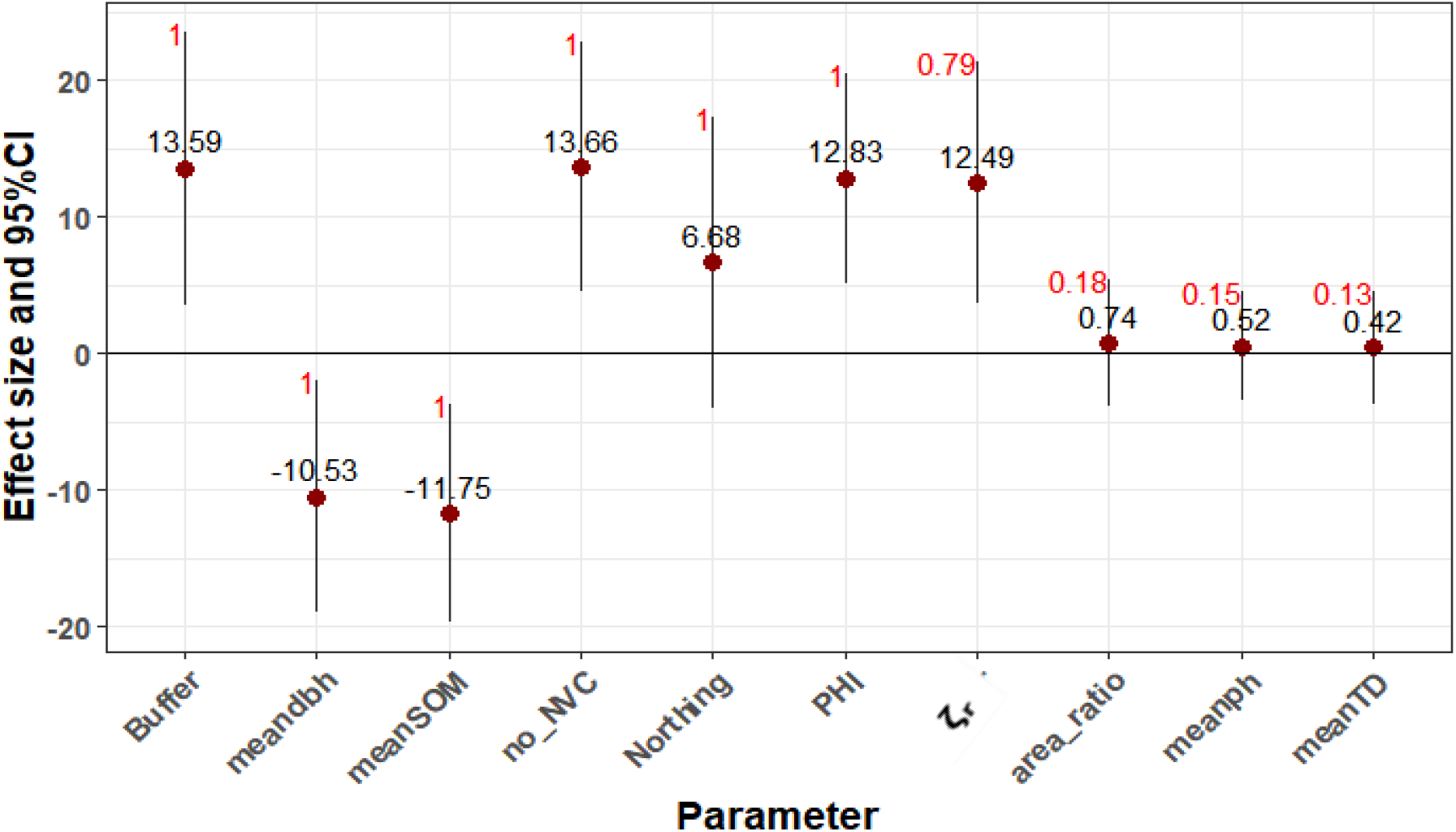
Model averaged effect sizes for woodland richness modelled with all variables using δ < 2 for the model set. Variable importance is in red. The points show the average effect size and the bars show the 95% confidence intervals.

Table 2 shows the results from the modelling using the GBM to find the variables which have the most influence on species richness. Parameters with relative influence of 10 or above have been selected as being important for predicting richness. ζ_r_ and Heterogeneity Index (HI) are again shown as having an impact on species richness. The GBM also selected mean LBA, mean pH and Northing.

Together, model-averaging and the GBM suggest that nine variables affect species richness of the woodlands, were woodland is defined as the 3200m^s^ covered by the area of the 16 plots. Three of these represent heterogeneity: ζr, number of NVC codes and HI. Of these, number of NVC codes and HI are present in every model in the model set and ζr is present in 80% of the models. The GBM selects ζ_r_ as the most influential variable.

### SCALE DEPENDENCE OF SPECIES RICHNESS

Figure 5 below shows the results of calculations to compute the exponent, z, at the two scales of this work.

**Figure 5.**
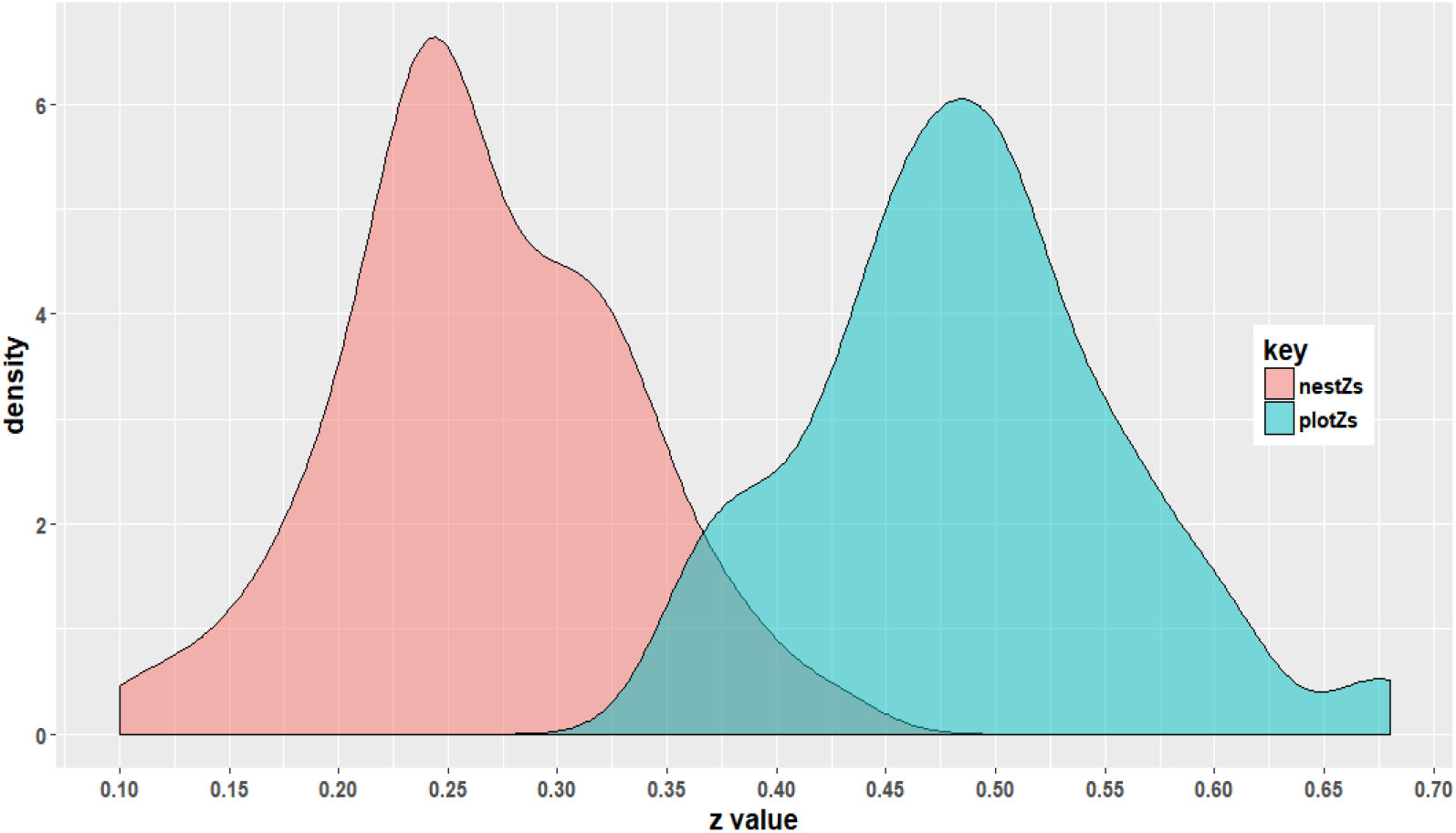
Distribution of z values at two scales. The pink distribution shows within-plot scale – 4m^2^ to 200m ^2^. The blue distribution shows between-plot scale – 200m^2^ to 3200m^2^

At the smaller within-plot scale the exponents are distributed around 0.25, whilst at the larger, between-plot scale they are distributed around 0.5.

Figure 6 shows the model-average effect sizes for (within-plot) nest z. The site level heterogeneity parameters HI and ζ_r_ have a positive effect on the value of z while mean SOM has a negative effect. The mean SOM is contained in every model in the model set, but the probability of inclusion of HI is 82% and for ζ_r_ only 57%, this implies that ζ_r_ is not an important parameter at nest scale. HI is the most significant parameter, the effect size double that of mean SOM.

**Figure 6.**
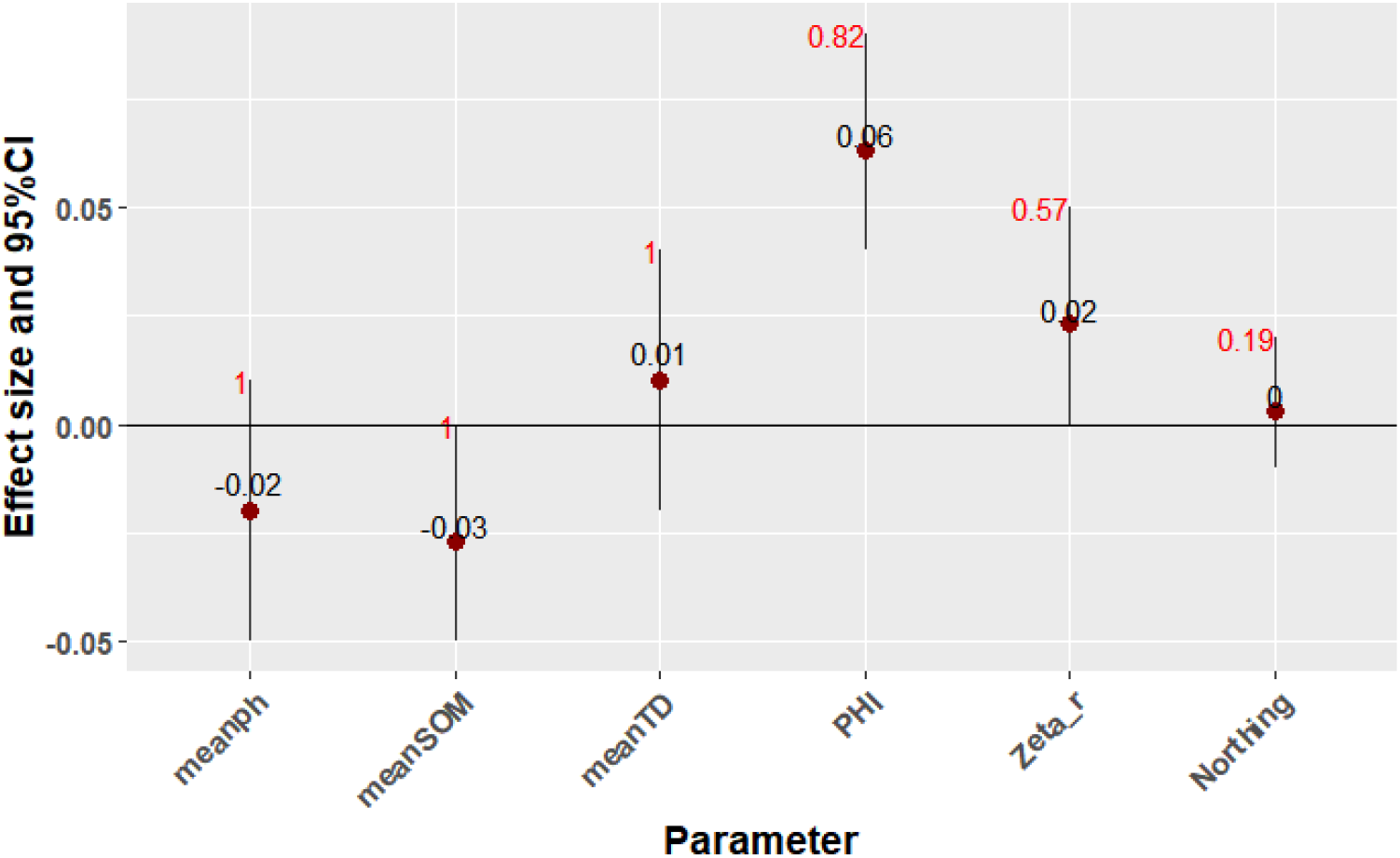
Model averaged effect sizes for nest z modelled with all variables using δ < 1.5 for the model set.

Figure 7 shows the model-average effect sizes for the slope of the between-plot specie area curve z. ζ_r_ is highly significant and has a positive effect on plot z. At this larger scale the effect of ζ_r_, which quantifies species turnover across the woodland, has much greater significance than at the smaller nest scale. Mean tree density and mean SOM are also significant, although less so than ζ_r_ and their effect sizes are less than a third of ζ_r_. Both mean SOM and mean tree density have a negative effect on plot z.

**Figure 7.**
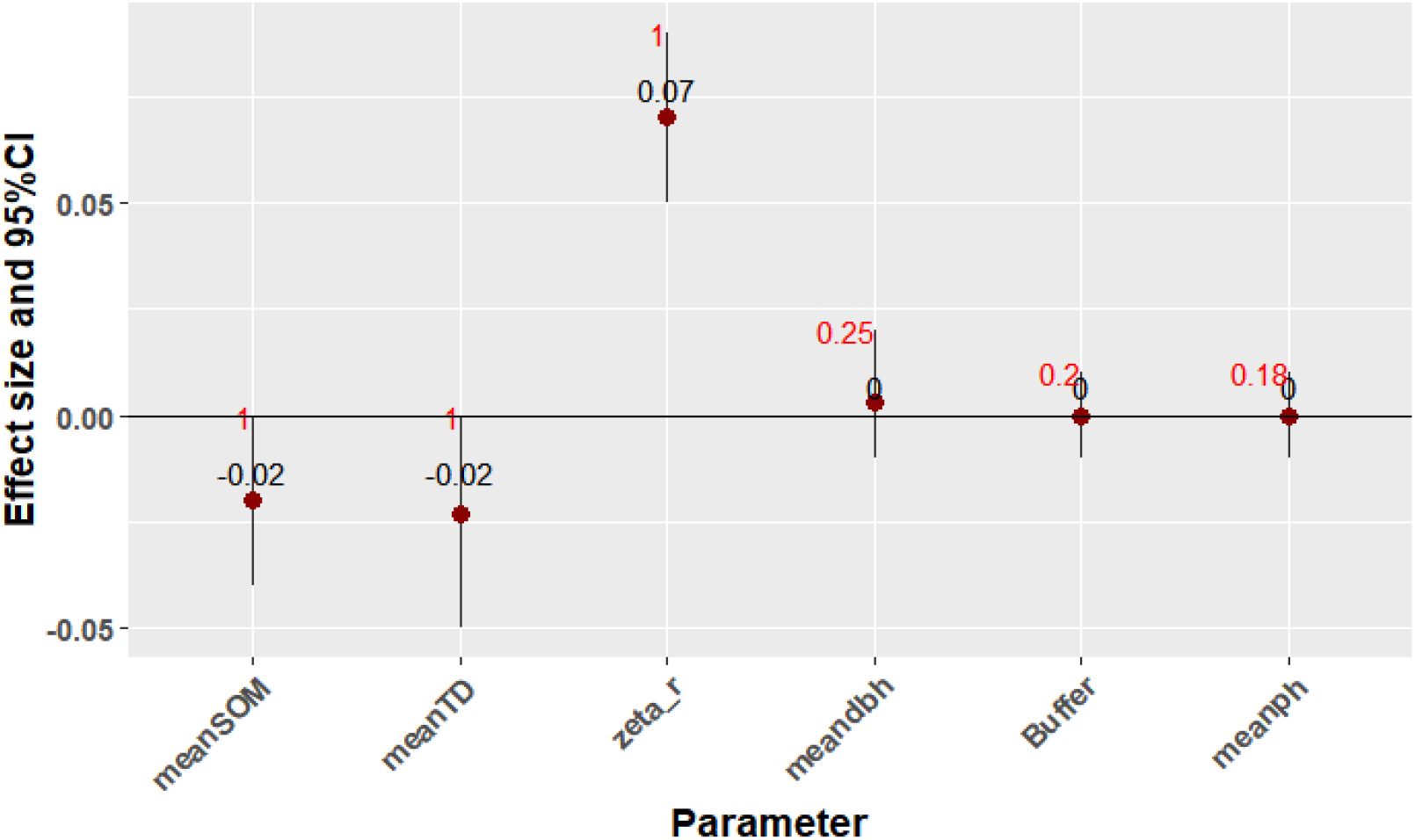
Model averaged effect sizes for plot z modelled with all variables using δ < 1.5 for the model set.

## Discussion

### VARIABLES AFFECTING SPECIES RICHNESS

We would expect that a suite of factors significant influence α, β and λ diversity in each woodland site. Our results indicate that the strongest effect is attributable to the heterogeneity of abiotic conditions at plot and site scale. We discuss the importance of heterogeneity and other influences below.

### HETEROGENEITY

The greater the HI, the more this indicates evidence of woodland management, open areas and riparian zones. This implies that all these features are associated with elevated plant species richness. The number of NVC codes and ζr quantify the number of different species assemblages. Different species assemblages are more likely to occur if environmental conditions differ across the woodland – for example, different levels of soil moisture or different light levels, again stressing the importance of heterogeneity within the woodlands. Moreover, it appears that increased site heterogeneity not only leads to increased gamma diversity, as seen by increased site species richness, but also increased beta diversity, as seen by the dependence of nest z on HI.

### MEAN-DBH AND MEAN-LBA

Model-averaging shows mean DBH to have a negative effect on richness and the GBM selected mean LBA as influential. This is to be expected, as an increase in either variable implies succession in the woodland (Packham et al., 2001; Smart et al., 2014) which leads to increased homogeneity because of a strengthening of the shading filter on the understorey species pool (Keith et al., 2009), again indicating the importance of woodland management schemes as well as sufficient area for natural disturbance dynamics to vary the successional status of the woodland.

### AREA OF SEMI-NATURAL HABITATS AROUND THE SITE (BUFFER)

The amount of semi-natural habitats around the woodlands (buffer) had a significant positive effect on richness and occurred in every model in the model set. The size of this buffer relates to the amount of the woodland that is exposed to natural habitats. The greater that value, the more protected the woodland is from agrochemicals and airborne environmental pollutants; particularly around the edges (Draaijers, et al., 1988), which are linked to reduced species richness (Bobbink et al., 1998; Pallet et al., 2016, Simkin et al., 2016; Stevens et al., 2016). Previous research has suggested that buffer regions should be included around woodlands for various reasons, including protection from anthropogenic influences (McWilliam et al., 2010) and vehicle pollution (Bignal et al., 2008). A larger the buffer may also imply a more connected the woodland; habitat fragmentation is known to have a negative impact on biodiversity, (Quine, 2011, Watts et al 2008). It is not possible within this dataset to ascertain the mechanism for the increase in richness with buffer, which might not be metapopulation buffering and reduction of extinction probability but could be because a high amount of seminatural habitat is a surrogate for the woodland being on a substrate and in a climatic space that favours a high gamma diversity.

### NORTHING

Northing is selected as an influential variable by the GBM. This would appear to be in contradiction to previous authors, (Gilman et al., 2014; Ohlemuller & Wilson, 2000, Gaston, K.J, 2000) who suggest that species richness would be expected decrease with latitude. However, the large-scale effects which are behind the generally accepted increase in species richness toward the equator are unlikely to be seen across the small change in latitude of our study, and also are likely to be dominated by other factors. For example, a bivariate plot of Northing with mean TD (figure 8) shows a negative correlation, suggesting that the more northerly woods are less dense and hence the increased richness could be due to the addition of more light-demanding species.

**Figure 8.**
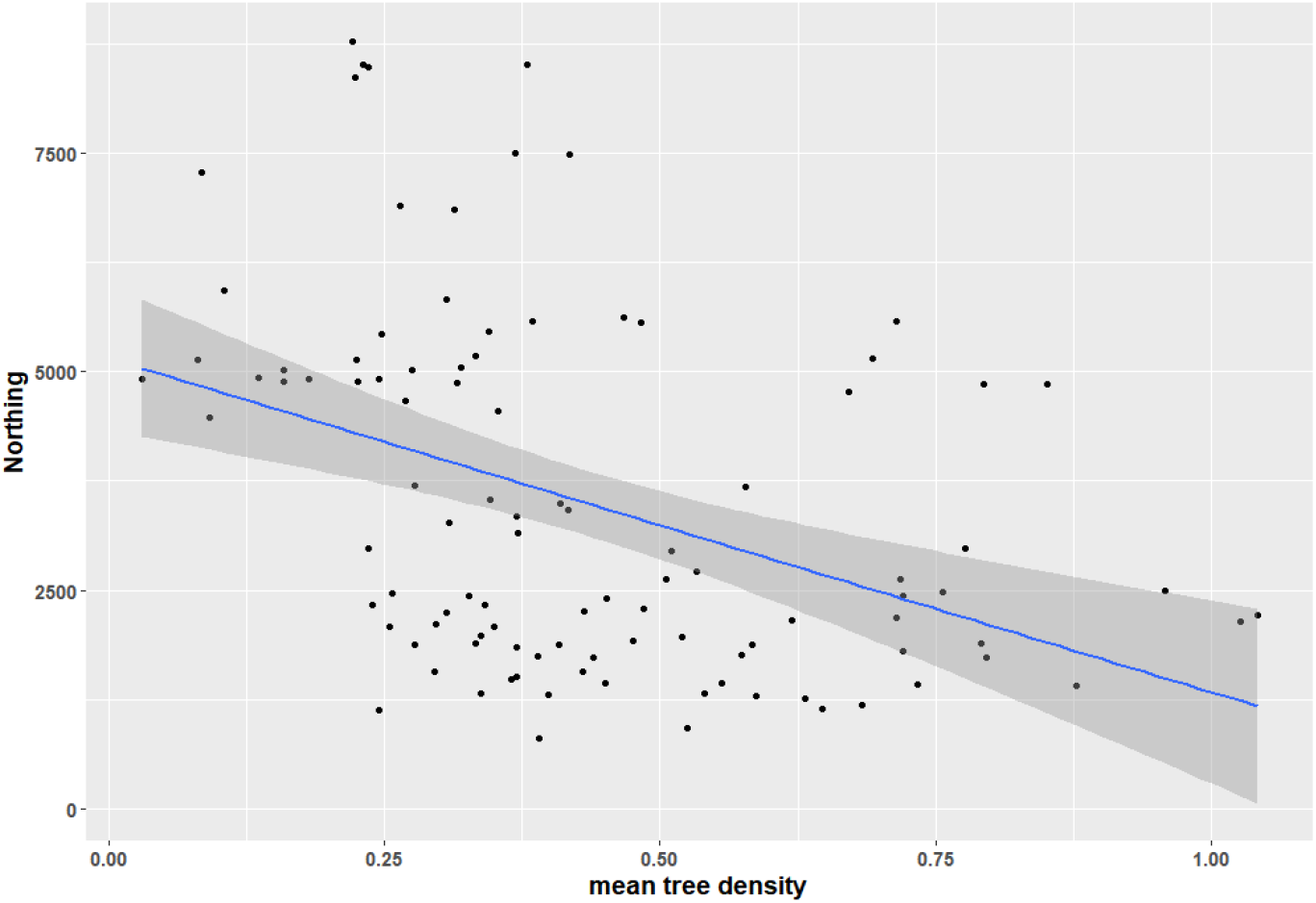
Scatter plot of northing with mean tree density suggests that the northern woodlands are more open. This could allow the inclusion of more light-loving species assemblages and hence add to species richness.

The Scottish woodlands might also consist of plant communities which tend to be richer. For example, if Scottish woodlands were predominantly W6 eutrophic Alder woodlands, they would be richer than W14 beech woods which are common in the South. However, figure 9 shows that this is not the case. The distribution of NVC communities is similar in the North and South. What is noticeable, however, is that for the same community, richness can be greater in Scottish woodlands, (figure 10). This suggests that other variables, not availabe in this dataset, may be responsible for the increased richness in Scottish woodlands, or that a combination of positive predictors is more likely to co-occur in the more Northern woodlands.

**Figure 9.**
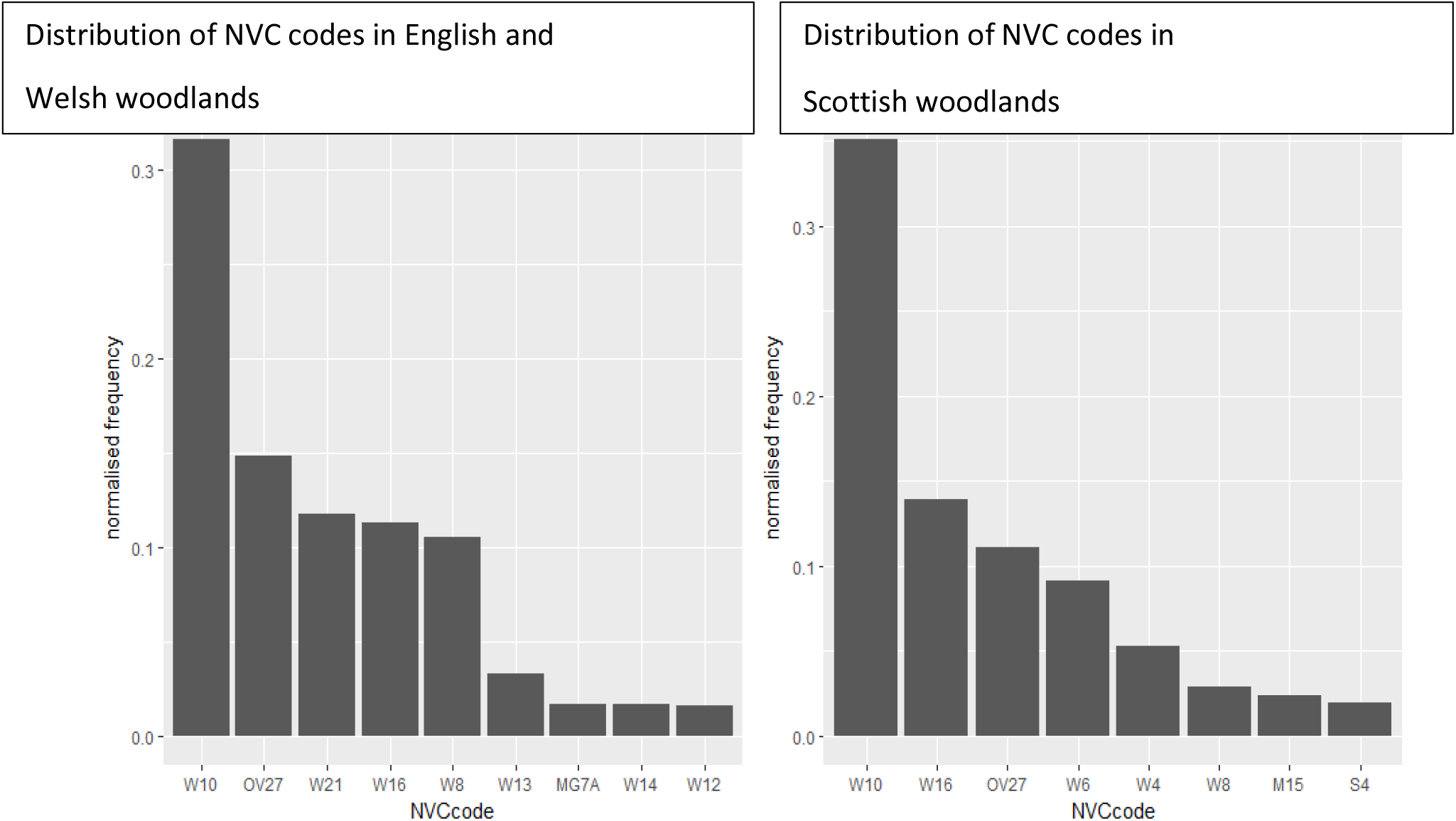
Distribution of NVC codes in Scotland and England and Wales. In both areas 30% of communities are W10, Quercus robur woodlands. These communities have one of the widest range of species richnesses of all NVC communities in this dataset.

**Figure 10.**
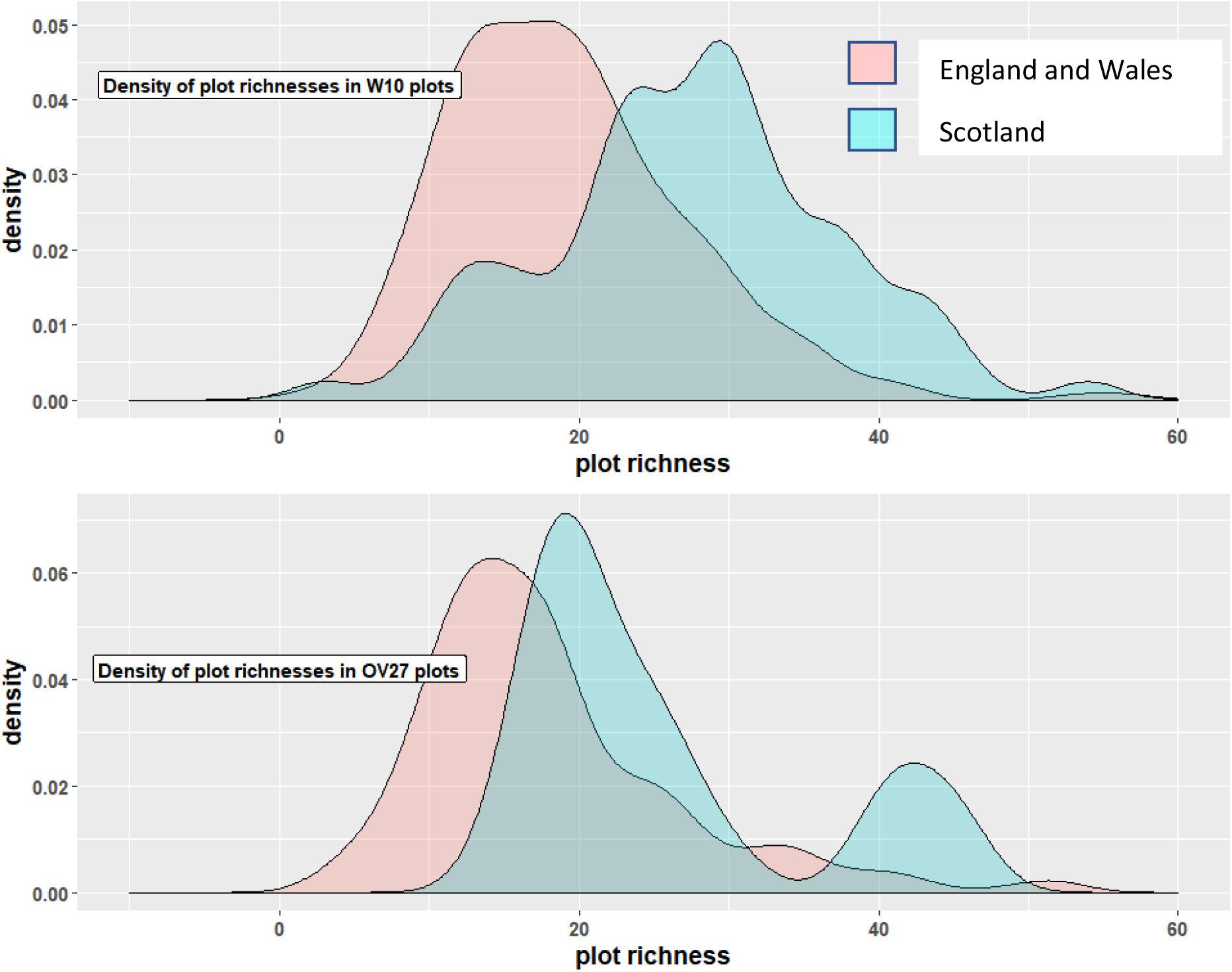
Density of plot richness for the most common NVC codes in the dataset in Scotland and England and Wales. Given the same NVC community, richness tends to be greater in Scotland for these common communities. Although OV27 is not a woodland community, it was was common in this dataset, demonstrating open areas within the wood.

### SOIL ORGANIC MATTER

SOM content was shown to have a significant and important negative effect on species richness in agreement with Koojiman & Cammeraat (2010). Organic soils have low pH and low pH soils in woodlands results in low species richness, (Cornwell & Grubb, 2003). However, in this data, very few of the plots were located on organic soils. The six NVC codes shown in table 4 represent 82% of all plots surveyed and give an indication of the majority of soil types.

**Table 4.** Soil types for the most common NVC codes in this data

| NVC code | % of plots | Soil type |
| --- | --- | --- |
| W6<br>*Alnus glutinosa-Urtica dioica* | 5% | Eutrophic moist soil alluvial or on flood fen peat |
| W8<br>*Fraxinus excelsior – Acer campestre – Mercurialis perennis* | 9.5% | Dry calcareous soils |
| W21<br>*Crataegus monogyna – Hedera helix* scrub | 10% | Various, base rich |
| W16<br>*Quercus* spp. – *Betula* spp. – *Deschampsia flexuosa* | 11.5% | Very acidic, oligotrophic, pH<4, sandy podzolic |
| OV27<br>*Epilobium angustifolium* Open habitat | 14% | Various, fertile, damp |
| W10<br>*Quercus robur – Pteridium aquilinum – Rubus fruticosus* | 32% | Neutral brown earths |

Of the most common NVC types in this dataset, only W6, 5% of the total, are potentially on peat, but W6 is a high nutrient, high richness community. It is therefore unlikely that the reduced richness with SOM is caused by reduced richness expected on organic soils.

The reduced richness could be due to low pH on other soil types since increased SOM has been shown to decrease pH (Russell, 1960; Williams & McDonald, 1960). On the other hand, adding SOM can also increase pH, depending on the initial acidity of the soil and the nature of the litter (Ritchie & Dolling, 1985). In this analysis, increased SOM appears to correlate with either decreased or no change in pH (figure 11).

**Figure 11.**
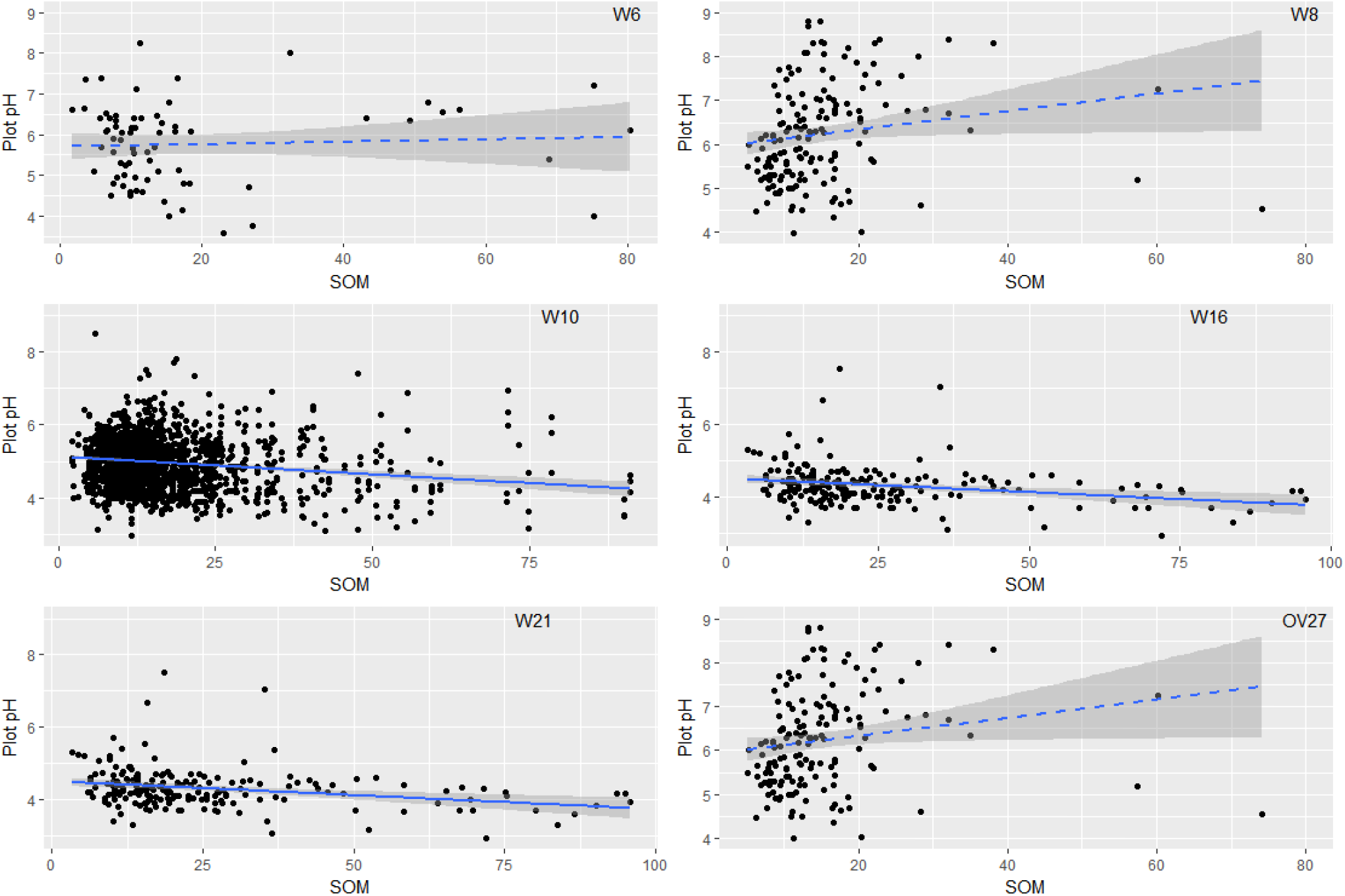
Change in plot pH with SOM content. For W10, W16 and W21 plots the pH reduces with increased SOM but this effect is not clear for other plot types

Figure 12 shows that all the common NVC communities tend to show reduced richness as SOM increases, regardless of soil type. Given also that the NVC code represents a community with similar physical conditions, such as nutrient levels, soil pH and light levels, and while the response to richness with increased SOM is seen across all communities, this finding implies that the variation in richness with SOM is unlikely to be due solely to one of those physical conditions, but perhaps to another factor not measured in this dataset.

**Figure 12.**
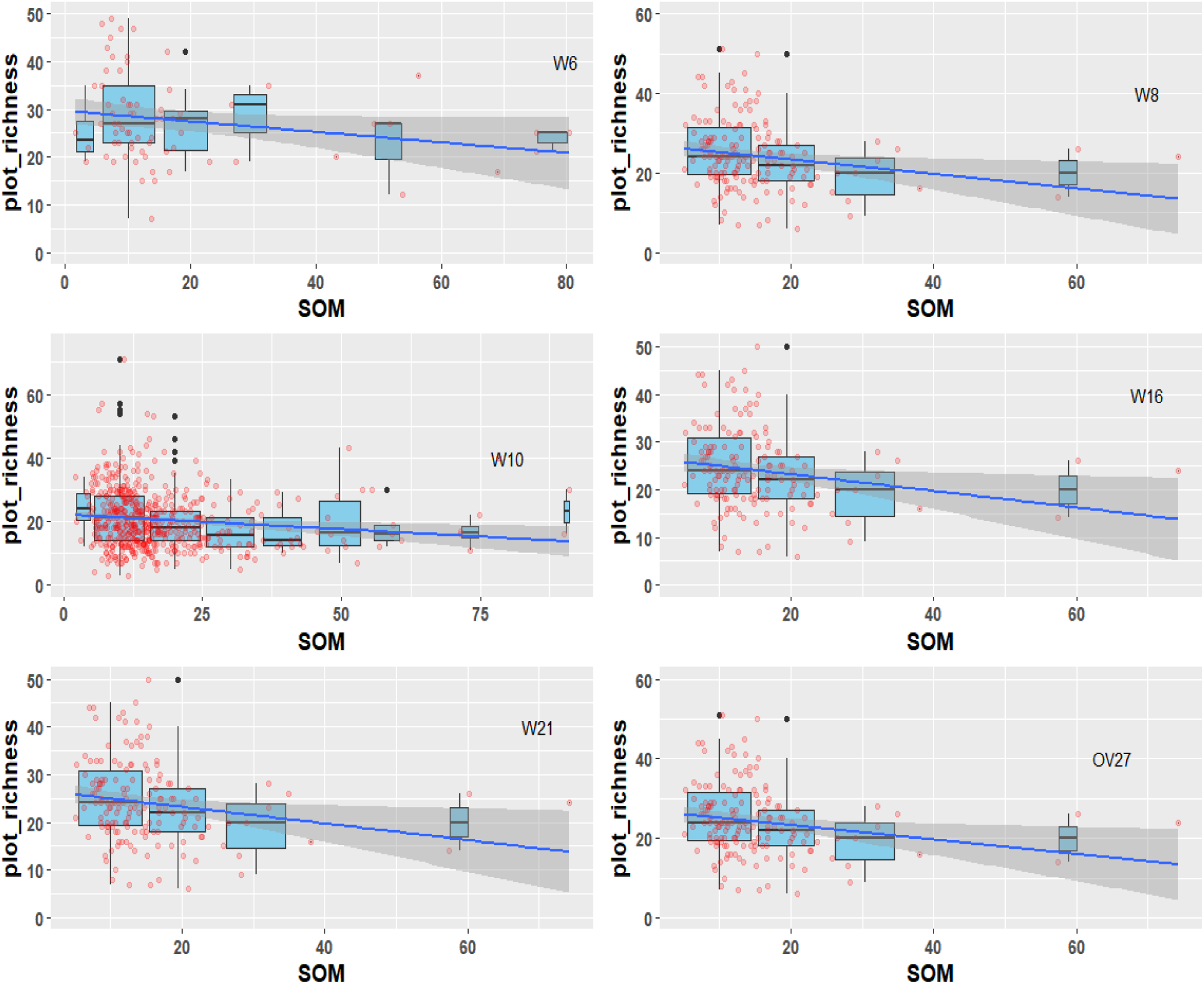
Plot richness with SOM for the six most common NVC communities in this data. The blue boxplots show the range in richness when grouped, the red dots are the raw ungrouped data. The variation in richness is high when SOM is low and vice versa.

Increased nitrogen deposition can both increase SOM (Fang et al., 2014; Zak et al., 2016) and reduce richness (Bobbink et al., 1998; Pitcairn et al., 2002; Simkin et al., 2016; Stevens et al., 2016). A smaller buffer could allow increased nitrogen deposition, (Draaijers et al., 1988; Kennedy & Pitman, 2004), which might be reflected in the Ellenberg N values (Maskell et al., 2010). Therefore, if woodlands with smaller buffers showed both increased Ellenberg N values and increased SOM, nitrogen deposition might be the mechanism behind the change in SOM and the reduced richness. In support of this figure 13 shows that buffer is negatively correlated with Ellenberg N values, but, in contradiction, it is positively correlated with SOM content. The latter association may be confused by the location of plots within the woodland.

**Figure 13.**
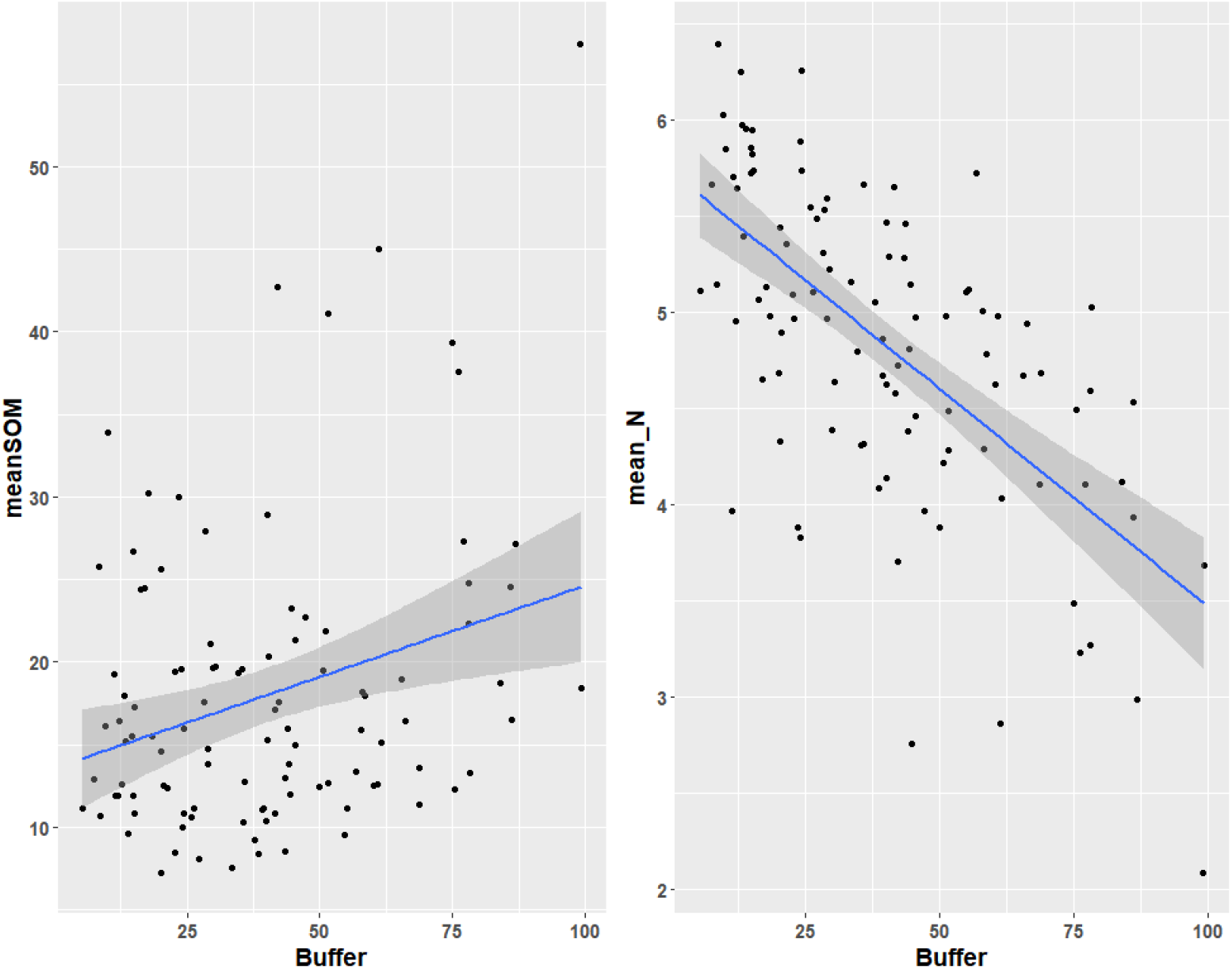
Change in mean SOM and Ellenberg N values with Buffer size. If nitrogen deposition was leading to increased SOM, then woodlands with smaller buffers would have greater SOM and larger Ellenberg N. The graph on the right shows that smaller buffers correlates with higher Ellenberg N, but not to greater mean SOM.

In summary, whilst SOM is negatively correlated with species richness, the causal mechanism is not evident. However, other authors, using modelled estimates of nitrogen deposition rather than Ellenberg N values with this dataset have shown positive correlation between SOM and modelled nitrogen deposition, (Kirby, 2005). It is also interesting to note that nitrogen deposition values were on average lower in Scotland (RoTaP, 2012), which may be another factor contributing to the increased richness of the more Northern woodlands.

### SOIL pH

The GBM selected mean pH as important for predicting species richness. Figure 14 demonstrates that in this data plot richness does have a unimodal response to plot pH, in agreement with Cornwell & Grubb, (2003)

**Figure 14.**
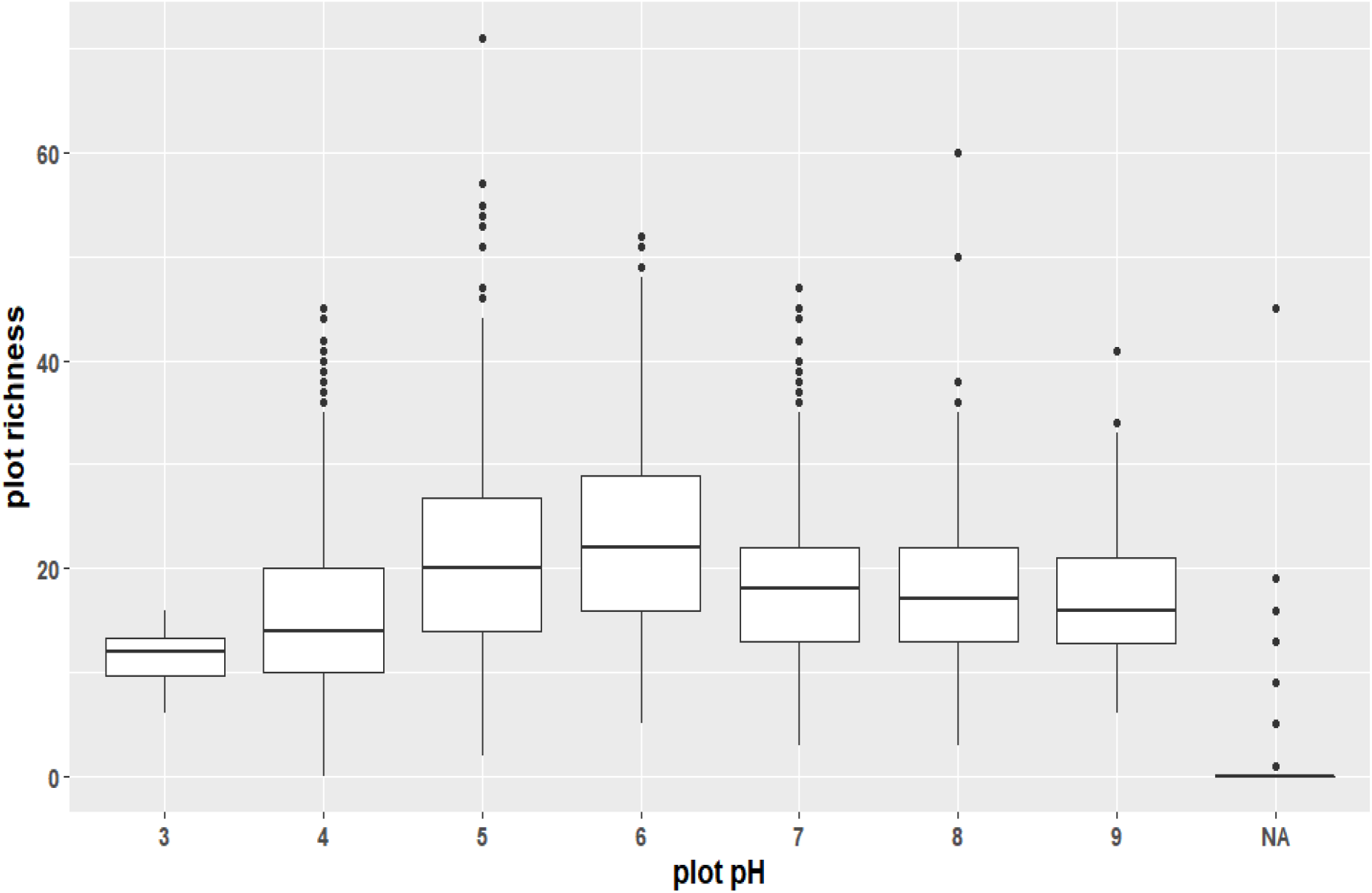
Plot richness response to pH can be seen to be unimodal with a peak around pH6.

### EFFECT OF DRIVERS OF SPECIES RICHNESS ON GRADIENT OF SPECIES AREA CURVE AT TWO SCALES

The fitted nest *z* values were distributed around 0.25. This agrees with the value of z predicted by other authors (Preston, 1962, Connor et al., 1983; Sugihara, 1976). At the larger scale, plot z was found to be distributed around 0.5. These two values correspond to those found in a similar study at two different scales by Crawley & Harral (2001) for vascular plants in the county of Berkshire, South East England. These authors comment that the smaller z value is expected at square meter scales, whereas the larger z value reflects large changes in species richness as new habitats are added when species are accumulated at larger scales.

At smaller scales, HI has a significant positive effect on nest z while SOM has a negative effect. This model suggests that biological variables appear to be influencing the slope of the species area curves, in contradiction to the remarks of other authors (Connor et al., 1983; Sugihara, 1976; Tjorve & Tjorve, 2017), but this is not necessarily the case. Both Sugihara (1979) and Tjørve & Tjørve (2017) point to the role of community structure and aggregation in shaping the species area curve. HI, by introducing elements into the woodland that increase plant species richness, is likely, on average, to be increasing the richness of each plot. This will imply a more diverse community over which the curve is constructed. Similarly, SOM can have a homogenizing effect on the plots due to loss of species richness, either directly (Kooijmann & Cammeraat, 2010) or because increased SOM indicates increased nitrogen deposition and eutrophication that favours the abundance of competitive species potentially suppressing species diversity in the understorey, (Maskell et al, 2010, Pallet et al 2016). Both HI and SOM can be seen therefore, as influencing community structure either positively or negatively and so effecting nest z. It should also be noted that model-averaged R^2^ for nest z was only 0.31, and that of the full model 0.33 so neither of the variables explained substantial variance in nest z.

At larger scales ζ_r_ has a significant positive effect on plot z, and mean SOM and mean TD have a significant negative effect. ζ_r_ is significant at the larger scale because it represents species turnover across the woodlands. The greater the species turnover, the greater the expected slope. Similarly, increasing mean TD and mean SOM are likely to have a homogenising effect on the woodland and therefore, to reduce plot z. At the larger scale, it is species turnover and heterogeneity that has been shown to influence the exponent of the species accumulation curves rather than biological factors. At this scale the model was more successful at explaining the variance in plot z, having R^2^ of 0.44 for the model-averaged model and 0.47 for the full model.

### VARIABLES NOT INCLUDED IN THIS ANALYSIS

Statistical modelling using model averaging explained 47% of the variance in plant richness. Since we chose only those variables that we both knew were likely to affect richness and that were available within the dataset, several important potential variables were omitted which may improve the explanatory power of the model. Woodland age and past land use have been shown to affect species composition (Hermy, 1994) and richness (Dzwonko & Loster, 1989; Peterken & Game 1984). Ancient woodlands are more likely to contain long-lived, poorly dispersing perennials whose migration to new woodland sites is limited (Kimberley, Blackburn & Whyatt, 2015). This constrains the potential richness of new woodlands, particularly if they are isolated. Ancient cultivation has also been shown to induce a species compositional gradient and thereby increase richness (Dupouey et al., 2002).

Plant available phosphorus has been shown to reduce species richness of native plants in alder forests (Hrivnak et al., 2015). Phosphorus eutrophication favours competitive species particularly at higher light levels and this can result in the exclusion of other species (Keersmaeker et al., 2004).

The amount of light reaching the woodland floor influences richness, with greater richness at higher light levels, (Kenedy & Pitman, 2004). However, eutrophication together with high light levels encourages shade-intolerant, competitive species which tend to reduce richness, (Cornwell & Grubb, 2003, Keersmaeker et al., 2004). Conversely, low pH in conjunction with low light levels can result in very little ground cover, (Cornwell & Grubb, 2003)

### USE OF ζ_r_ AS HETEROGENEITY METRIC

Results presented here suggest that ζr is a useful way to quantify species turnover. This variable was selected as the most influential by the GBM and was significant when used in model-averaging. The fact that ζr was a significant parameter when modelling the slope of species accumulation curves across the wood, but not when modelling the slope of species area curves across plots, demonstrates that it quantifies species turnover. This metric could be easily generated from species lists as an indicator to woodland managers of the number of different habitats within the woodlands.

### SYNTHESIS

Habitat heterogeneity was found to be important for species richness in UK broadleaved woodlands. Heterogeneity in these models included coppicing, riparian zones and open areas, suggesting that all these factors need to be considered if woodland plant diversity is to be increased. Isolation was negatively correlated with species richness and therefore any possibility to increase the size and connectivity of new woodlands should be a priority. The northern woodlands were found to be richer than those in the south, and more importantly, in some cases the same plant communities were richer in Scotland than the south. The reason for this was unclear and further work could be done, for example, examining interactions or nitrogen deposition data.

The parameter effect sizes for all the variables were similar, so that no one factor can be said to be most important for richness. This implies that in order to increase woodland richness all must be addressed simultaneously, and that one single factor will neither explain the species richness or lack of diversity in our woodlands.

